# Stochastic variation in the FOXM1 transcription program mediates replication stress tolerance

**DOI:** 10.1101/2024.03.26.585806

**Authors:** Hendrika A. Segeren, Kathryn A. Wierenga, Frank M. Riemers, Elsbeth A. van Liere, Bart Westendorp

## Abstract

Oncogene-induced replication stress (RS) is a vulnerability of cancer cells that forces reliance on the intra-S-phase checkpoint to ensure faithful genome duplication. Inhibitors of the crucial intra-S-phase checkpoint kinases ATR and CHK1 have been developed, but persistent proliferation and resistance to these drugs remain problematic. Understanding drug tolerance mechanisms is impeded by analysis of bulk samples, which neglect tumor heterogeneity and often fail to accurately interpret cell cycle-mediated resistance. Here, by combining intracellular immunostaining and RNA-sequencing of single cells, we characterized the transcriptomes of oncogenic RAS-expressing cells that exhibit variable levels of RS when challenged with a CHK1 inhibitor in combination with the chemotherapeutic drug gemcitabine. We identified 40 genes differentially expressed between tolerant and sensitive cells, including several FOXM1 target genes. While complete knockdown of *FOXM1* impeded cell proliferation, a partial knockdown protected cells against DNA damage, and improved recovery from drug-induced RS. Our results suggest that low levels of FOXM1 expression protects subsets of oncogenic RAS-expressing cells against DNA damage during drug-induced replication stress.

## Introduction

Oncogene-induced replication stress (RS) is a vulnerability of cancer cells that can be exploited by anti-cancer therapies. Seminal studies in the beginning of this century already showed that oncogenes, such as RAS, induce DNA damage in precancerous lesions (Bartkova et al., 2006, Gorgoulis et al., 2005). Further research revealed that oncogene-induced RS underlies the elevated levels of DNA damage, and that RS is present in the vast majority of human tumors. As a result, RS is proposed as an emerging hallmark of cancer (Macheret, Halazonetis, 2015b).

RS is defined as stalling of the replication fork, which can arise due to shortage of substrates, collisions between replication and transcription machinery, or DNA lesions or secondary structures that hinder the replication machinery. Unresolved RS can progress to replication fork collapse, resulting in single- and double-stranded DNA breaks. To prevent this, cells respond to RS by triggering the intra S-phase checkpoint. Briefly, this checkpoint is initiated when Replication Protein A (RPA) binds to single-stranded DNA that is exposed upon uncoupling of helicase and polymerase activity during fork stalling. This triggers recruitment and activation of ATR and its downstream kinase CHK1, which together induce a cascade of kinase activation that acts to stabilize and repair the stalled replication fork, fire dormant origins in the vicinity of the stalled fork, attenuate global DNA replication and slow down cell cycle progression. This multifaced response ensures faithful genome duplication before mitosis (Lecona, Fernandez-Capetillo, 2018).

In general, loss of ATR or CHK1 is lethal in cells where oncogenes are activated (Murga et al., 2011, Gilad et al., 2010, Oo et al., 2018, Schoppy et al., 2012). On the basis of this knowledge, inhibitors against key-players of the intra S-phase checkpoint are developed and currently evaluated in clinical trials (Baillie, Stirling, 2021). To potentiate the effect of intra S-phase checkpoint ablation, ATR and CHK1 inhibitors can be combined with a low dose of chemotherapeutic drugs (Liu et al., 2017, Wallez et al., 2018). However, drug resistance remains a major problem (Hong et al., 2018). The limited *in vivo* activity of drugs which exacerbate RS suggests that cancer cells employ strategies to tolerate RS. Indeed, stabilization of the replication fork (Bianco et al., 2019), increased expression of RPA (Bélanger et al., 2018), and increased dormant origin firing (Jo et al., 2021) grant RS tolerance. Interestingly, factors that curb RS in cancer cells, such as CLASPIN, CHK1, and NRF2, frequently display increased transcript levels in cancer cells (Bianco et al., 2019, Bertoli et al., 2016, Mukhopadhyay et al., 2020). Moreover, unbiased screening approaches uncovered that cell cycle related genes mediate resistance to intra S-phase checkpoint inhibitors (Blosser et al., 2020, Schleicher et al., 2020, Ruiz et al., 2016). However, since these studies employed bulk sample approaches, transcriptional heterogeneity was neglected, rare resistance-conferring events missed, and the role of cell cycle progression potentially misinterpreted. As a result, the development of novel clinical strategies based on these studies is rare.

The importance of single-cell data in drug-resistance studies is highlighted by Shaffer *et al*., who unveil that rare cancer cells express resistance genes prior to treatment to resist therapy (Shaffer et al., 2017). In support of this notion, treatment with RS-inducing drugs leads to a reduction in the number of transcriptionally distinct clones (Seth et al., 2019), suggesting selection pressure for cells harboring drug-tolerant characteristics. Besides pre-existing heterogeneity, it is becoming increasingly evident that cancer cells modulate their transcriptome upon treatment to circumvent therapy. For example, chemotherapeutic drugs may induce a transient drug-tolerant state in a subpopulation of cells (Goldman et al., 2015, Rehman et al., 2021). It is hypothesized that this provides a time window in which permanent resistant cells can arise. Because transcriptional heterogeneity could result in resistance to RS-inducing drugs and consequently tumor relapse, the mechanisms underlying RS tolerance warrant further investigation.

Here, we employed a strategy in which we combine immunostaining of RS markers and information on cell cycle phase with single cell RNA-sequencing. This allowed us to shed light on the biological variability in response to RS. We uncovered a subset of genes with an altered expression profile in cells that maintained low levels of RS despite challenge with RS-inducing drugs. We also identified genes that make cells more sensitive to replication stress, which included several FOXM1 target genes. Consistent with this, partial knockdown of FOXM1 mitigated DNA damage and improved cell survival following treatment with RS-inducing drugs. These findings provide potential new avenues for development of synthetic lethality strategies and identification of biomarkers to optimize anti-cancer therapy.

## Results

### γH2AX is a replication stress marker suitable for flow cytometry of DSP-fixed cells

To unmask transcription mediated RS-adaptation mechanisms, transcriptomic information and the level of RS in single cells needs to be combined. Therefore, we adapted a previously published strategy in which cells are reversibly fixed to allow antibody staining while preserving RNA for sequencing (Gerlach et al., 2019). As summarized in Fig. 1A, cells were fixed using the chemically reversible crosslinking reagent DSP. Next, cells were stained with an antibody that recognizes an RS-specific marker and sorted based on RS levels in 384-well plates using FACS. Subsequently, de-crosslinking was performed using the reducing agent DTT and cells were subjected to first strand cDNA synthesis and single-cell RNA-sequencing.

**Figure 1:**
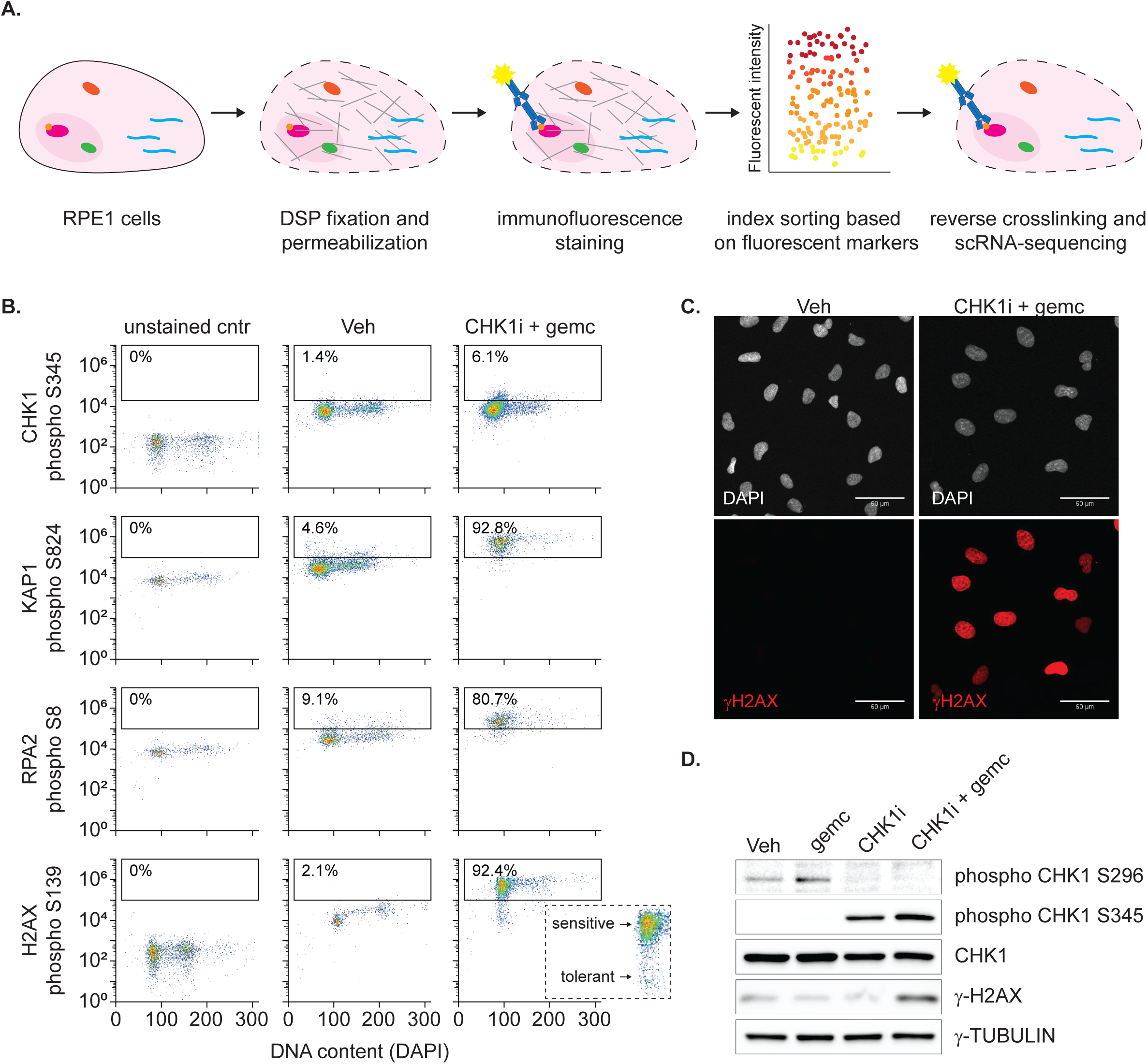
γH2AX is a replication stress marker suitable for flow cytometry of DSP-fixed cells. **A** Schematic overview of the technique to combine immunostaining and single-cell RNA sequencing. Cells are fixed with DSP, permeabilized and stained using fluorescent antibodies. Next, cells are sorted based on fluorescent intensity. After de-crosslinking, cells are subjected to single-cell RNA sequencing. **B** Flow cytometry data showing the intensity of potential RS markers in individual RPE-HRAS^G12V^ cells treated for 24 hours with 10 nM CHK1i + 100nM gemcitabine or vehicle (Veh). Unstained control refers to control cells not incubated or, when applicable, incubated with the secondary antibody only. Inset zooms in on cells stained for γH2AX after treatment with CHK1i + gemcitabine to indicate heterogeneity in γH2AX level. **C** Representative example of γH2AX immunostaining on RPE-HRAS^G12V^ cells treated for 24 hours with 10 nM CHK1i + 4 nM gemcitabine or vehicle (Veh). **D** Immunoblot showing synergistic induction of RS by 10 nM CHK1i + 4 nM gemcitabine in RPE-HRAS^G12V^ cells, as indicated by phospho CHK1 S345 and γH2AX. The absence of phosphorylation of CHK1 on its autophosphorylation site S296 indicates effective inhibition by CHK1i.

Before implementation of this technique, we first investigated which antibody against RS-induced protein modifications is compatible with DSP-fixation and analysis by flow-cytometry. To induce RS, we employed the frequently used chemotherapeutic drug gemcitabine in combination with the CHK1 inhibitor (CHK1i) prexasertib (Segeren et al., 2022). In response to RS, the intra S-phase checkpoint kinase ATR is activated to stabilize and repair stalled forks and delay cell cycle progression. This is mediated by a sequence of events including phosphorylation of CHK1, RPA2, KAP1 and H2AX (Toledo et al., 2013, Branigan et al., 2021). Antibodies against phosphorylated variants of these proteins are previously shown to detect increased levels of RS by flow-cytometry (Atashpaz et al., 2020, Branigan et al., 2021). We assessed if these antibodies could detect an increase in phosphorylated CHK1, RPA2, KAP1 and H2AX in DSP-fixed cells after treatment with CHK1i + gemcitabine.

We made use of RPE-1 cells harboring a doxycycline-inducible variant of oncogenic RAS (hereafter referred to as RPE-HRAS^G12V^ for cells with doxycycline-induced expression of HRAS^G12V^ or control for their non-induced counterparts). (Segeren et al., 2022). The advantage of this system is that adaptation to RS can be studied in the frequently occurring oncogenic context of RAS hyperactivation, while the effect of other tumor-specific mutations is excluded. We previously described that RPE-HRAS^G12V^ cells show mild endogenous RS and markedly enhanced sensitivity to CHK1i + gemcitabine (Segeren et al., 2022)). However, control RPE cells also show RS in presence of high doses of these drugs. Accordingly, treating RPE control cells with a high dose of CHK1i + gemcitabine resulted in 2N-cell cycle arrest, as seen by the accumulation of cells with low DAPI signal, indicating severe stress. CHK1i + gemcitabine also triggered an abundant increase in phosphorylated KAP1, RPA2 and H2AX (Fig. 1B). However, the tested antibody against phospho-Serine 345 on CHK1 failed to show an increase in this flow cytometric analysis of DSP fixed cells, excluding it as an RS-marker for this project (Fig. 1B). While KAP1 mediates RS-induced DNA remodeling and RPA2 protects stalled replication forks, phosphorylated H2AX is present at collapsed replication forks (Goodarzi, Jeggo & Kurka, 2011, Toledo et al., 2013). Since the latter is the most downstream event in the RS-cascade and indicates severe RS, the antibody against phosphorylated H2AX Serine 139 (hereafter referred to as γH2AX) was selected as a proxy for RS-induced DNA damage. Interestingly, flow-cytometry analysis of γH2AX stained cells revealed great diversity in the signal, and presumable RS-level, between individual cells (Fig. 1B, inset). Consistent with this, heterogenous phosphorylation of H2AX S139 in response to RS was confirmed by immunofluorescence staining (Fig. 1C). In addition, immunoblotting confirmed that substantial γH2AX was observed when the CHK1i prexasertib was combined with a low dose of gemcitabine, but not with either drug alone (Fig. 1D). Based on these observations, we concluded that the antibody against γH2AX can be used to determine the level of RS induced by CHK1i and gemcitabine at a single cell resolution in DSP-fixed cells.

### High quality single-cell RNA sequencing data of fixed cells with known level of replication stress

After identification of γH2AX as an RS-marker we directly compared fresh and DSP fixed cells to evaluate the extent to which fixation with DSP affects the quality of single-cell RNA sequencing data. Since the response to RS is affected by cell cycle stage, we decided to sort only cycling cells. To this end we made use of the fact that our RPE-HRAS^G12V^ cells stably expressed the Fluorescent Ubiquitination-based Cell Cycle Indicator (FUCCI4) system (Bajar et al., 2016). We sorted RPE-HRAS^G12V^ cells expressing Geminin_1-110_, representing S/G2-phase, with and without treatment with RS-inducing drugs. Half of the cells were directly sorted (fresh), whereas the other half was first fixed with DSP and stained using the aforementioned γH2AX antibody. All samples were subjected to standard cDNA preparation, including de-crosslinking, and RNA-sequencing. After initial quality control (described in methods section), 232 fresh (success rate = 60.42%) and 273 DSP-fixed (success rate = 71.09%) cells were selected for downstream analysis. The mean number of identified genes (5644 in fresh versus 5044 in DSP-fixed cells) was comparable (Fig. 2A), as was RNA count, percentage of mitochondrial genes, and spike-in RNAs (Fig. S1A-C). Moreover, the average gene expression and gene detection rate were not affected by DSP-fixation (Fig. 2B-C). In addition, the similar coefficients of variation in the two cell populations indicates that DSP-fixation does not negatively impact the ability to detect expression heterogeneity (Fig. 2D).

**Figure 2:**
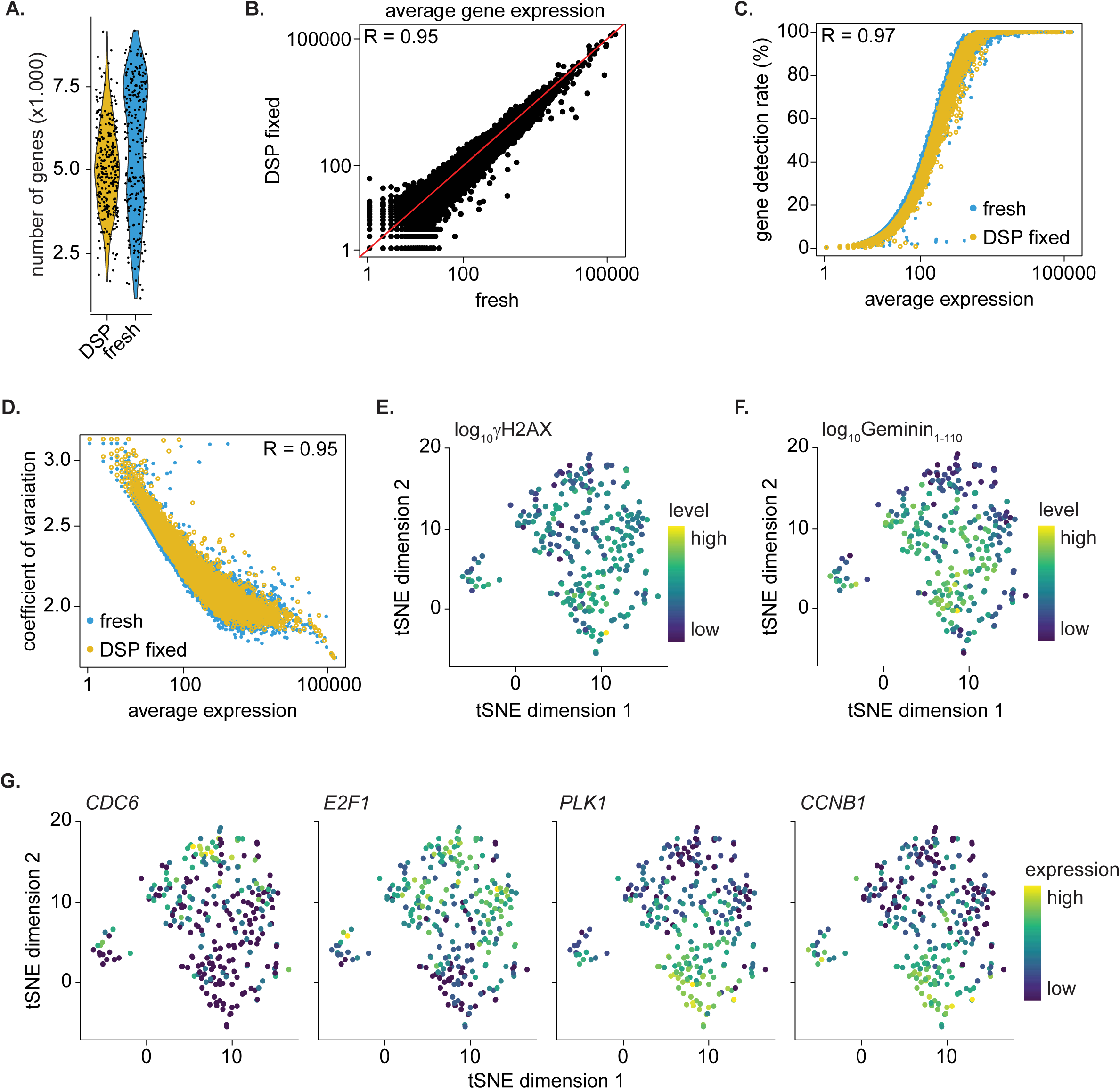
High quality single-cell RNA sequencing data of fixed cells with known level of replication stress. **A** Violin plot representing the average numbers of genes detected per cell in fresh and DSP fixed RPE-HRAS^G12V^ cells. **B** Scatter plot showing the average gene expression in DSP fixed and fresh cells. R value indicates Pearson Correlation. The red line indicates x=y. **C** Scatter plot showing the correlation between the gene detection rate and average gene expression in fresh and DSP fixed cells. R value indicates Pearson Correlation coefficient between fresh and DSP fixed cells. **D** Scatter plot showing the correlation between the coefficient of variation and average gene expression in fresh and DSP fixed cells. R value indicates Pearson Correlation coefficient between fresh and DSP fixed cells. **E** Dimensionality reduction using tSNE of DSP-fixed cells. Cells are color coded according to γH2AX signal. **F** Feature plot in which cells on tSNE plot in E are color-coded according to mAG-Geminin_1-110_ signal. **G** Feature plots in which cells on tSNE plot in E are color coded according to the expression of S-phase (*CDC6* and *E2F1*) or G2-phase (*PLK1* and *CCNB1*) markers.

After confirming that we can obtain high quality single cell RNA sequencing data from DSP-fixed cells, we aimed to identify gene-expression programs that mediate the low level of RS in a subset of cells and potentially underly drug resistance. However, when we analyzed cells treated with Chk1i + gemcitabine, the γH2AX positive and negative cells do not clearly cluster apart in the tSNE plot shown in Fig. 2E. Thus, not the level of RS, but other factors account for the clustering within the DSP-fixed cell population. To assess if cell cycle status can explain the clustering, we plotted the protein level of the FUCCI4 cell cycle marker Geminin_1-110_. The protein level of Geminin_1-110_, which gradually increases during S-phase progression, correlated well with the different cell clusters, suggesting that transcriptional events underlying S and G2 phase account for clustering (Fig. 2F). Accordingly, the expression of early S-phase (*CDC6* and *E2F1*) and late G2/M-phase (*PLK1* and *CCNB1*) markers showed that high levels of RS are predominantly present in cells in late S or G2 phase (Fig. 2G). Thus, cell cycle position can be a major confounding factor when evaluating the transcriptomic response to RS.

### Identification of putative genes that confer replication stress tolerance

To reduce the variation in level of RS caused by cell cycle status, we more stringently selected cells solely in mid S-phase based on the DNA content using DAPI (Fig. 3A). Subsequently, we selected S-phase cells negative for γH2AX, and S-phase cells with low, medium or high levels of γH2AX using flow cytometry before and 16h after treatment with RS-inducing drugs. As seen previously (Fig. 1C), treatment with CHK1i + gemcitabine increased the level of γH2AX in RPE-HRAS^G12V^ cells, but several cells still maintained low levels of γH2AX (Fig. 3A). Hence, to allow identification of mechanisms that facilitate resistance to RS-inducing drugs, we collected cells with no, low, intermediate or high levels of γH2AX staining for single-cell RNA-sequencing.

**Figure 3:**
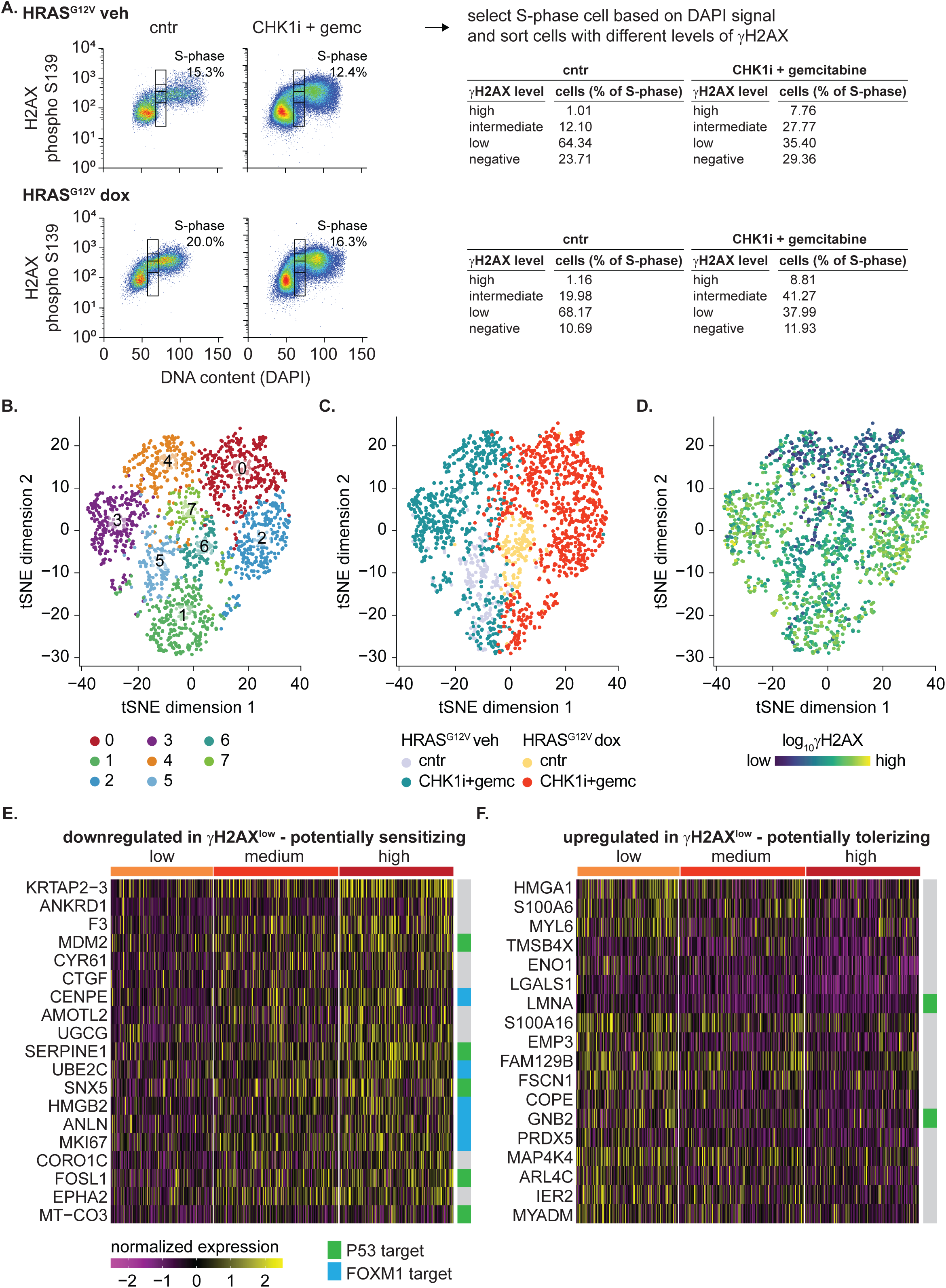
Identification of putative genes that correlate with replication stress tolerance. **A** Flow cytometry data of RPE-HRAS^G12V^ and control cells treated for 16 hours with 10 nM CHK1i + 4 nM gemcitabine or vehicle (Veh). Sorting strategy is shown: first S-phase cells were selected based on DAPI signal. For drug treated cells equal number of cells with no, low, medium or high levels of γH2AX were sorted. The percentages of cells in these different categories before sorting are indicated and show an increase in the cell population with high level of RS after treatment with CHK1i + gemcitabine. **B** Dimensionality reduction using tSNE of all cells (HRAS^G12V^ and control) before and after treatment with 10 nM CHK1i + 4 nM gemcitabine shows separate clusters of cells. **C** Feature plot in which cells on tSNE plot in B are color coded based on the different conditions. **D** Feature plot in which cells on tSNE plot in B are color coded according to γH2AX signal. **E** Heatmap of genes differentially expressed and downregulated in RPE HRAS^G12V^ γH2AX^low^ versus γH2AX^high^ control RPE cells 16 hours after treatment with 10 nM CHK1i + 4 nM gemcitabine. **F** Heatmap of genes differentially expressed and upregulated in γH2AX^low^ versus γH2AX^high^ RPE HRAS^G12V^ cells 16 hours after treatment with 10 nM CHK1i + 4 nM gemcitabine.

In an attempt to identify the influence of oncogenic RAS on transcriptional mechanisms of resistance, we similarly treated RPE-HRAS^G12V^ and control RPE cells with CHK1i + gemcitabine and sorted cells with different levels of RS. We subjected these cells to single-cell RNA sequencing and selected cells with more than 1000 unique RNA counts and expressing at least 500 genes for downstream analysis (exact parameters stated in Methods section). Next, we performed principal component analysis and t-SNE visualization. This revealed eight distinct clusters of cells that correlated well with the different experimental conditions (Fig. 3B,C). Interestingly, rare cells treated with CHK1i + gemcitabine are located within the untreated cell cluster (Fig. 3C), potentially representing non-damaged, RS-tolerant cells. Moreover, CHK1i + gemcitabine treated cells in cluster 0 and 4 display lower levels of RS compared to cells in cluster 2 and 3 (Fig. 3A-D).

To rule out the influence of cell cycle position, we compared the DAPI signal, indicative of S-phase progression, in cells with different levels of γH2AX signal. The DAPI signal was comparable in cells with low, medium, and high levels of RS, but the DAPI signal was much lower in γH2AX^negative^ cells (Fig. S2A). Because we suspected that the absence of RS in γH2AX^negative^ cells could be attributed to their earlier position in S-phase, we chose to compare the transcriptomes of γH2AX^low^ and γH2AX^high^ RPE-HRAS^G12V^ cells. We suspect that cells able to withstand DNA damage during replication stress represent cells within in a tumor that could survive treatment with RS-inducing drugs. Differential expression analysis revealed 19 genes that were significantly downregulated in γH2AX^low^ RPE-HRAS^G12V^ cells, suggesting that elevated levels of these genes are correlated with sensitivity to RS-inducing drugs (Fig. 3E and Table S1). A large subset of these genes (*CENPE*, *UBE2C*, *HMGB2*, *ANLN* and *MKI67)* are controlled by the key G2/M transcription factor FOXM1 (Fischer, Martin et al., 2016). In contrast to genes with a reduced expression in γH2AX^low^ cells, 18 genes, including several P53 target genes, had significantly higher expression in γH2AX^low^ cells, and thus correlated with RS tolerance (Fig. 3F and Table S1).

Next, we evaluated if the genes differentially expressed in γH2AX^high^ versus γH2AX^low^ cells are co-expressed (Fig. S2B). Among genes downregulated in γH2AX^low^ cells, the expression of *ANLN*, *HMGB2*, *CENPE*, *MKI67* and *UBE2C* correlated, which is expected as they are all regulated by the FOXM1 transcription factor. However, no co-expression of the putative RS-tolerance conferring genes, genes upregulated in γH2AX^low^ cells, was observed. This indicates that these genes are regulated independently of each other.

Notably, several FOXM1-target genes were also found to be downregulated in the γH2AX^low^ RPE control cells that lack expression of oncogenic RAS (Fig. S2C). This suggests that reduced activation of this transcriptional program in cells with decreased γH2AX levels is a general phenomenon and not necessarily linked to oncogene expression.

Altogether, these data indicate that a subset of oncogenic RAS expressing cells is protected from RS upon treatment with RS-inducing drugs and that these cells transcriptionally diverge from drug-sensitive cells, with many differentially expressed genes targeted by the transcription factor FOXM1.

### Validation of putative RS tolerance mechanisms

Next, we assessed if the aforementioned genes that were differentially expressed in γH2AX^low^ versus γH2AX^high^ RPE-HRAS^G12V^ cells could be functionally responsible for RS sensitivity and RS tolerance (Fig. 4A). To this end we knocked down these genes individually prior to treatment with CHK1i + gemcitabine and analyzed if this affected RS. We hypothesized that knocking down sensitizing genes would result in a decrease in replication stress while knocking down tolerizing genes would result in an increase in RS upon treatment with CHK1i+gemcitabine (Fig. 4A). We excluded P53 target genes (*MDM2*, *SERPINE1*, *SNX5*, *FOSL1*, *MT-CO3*) as the key role of P53 in the RS response is well-established (Macheret, Halazonetis, 2015a). Moreover, individual FOXM1 target genes (*CENPE*, *UBE2C*, *HMGB2*, *ANLN*, *MKI67*) were excluded from further analysis and replaced by knockdown of *FOXM1* itself to address the role of this entire transcription program in the RS-response (Fischer, M., 2017, Fischer, Martin et al., 2016).

**Figure 4:**
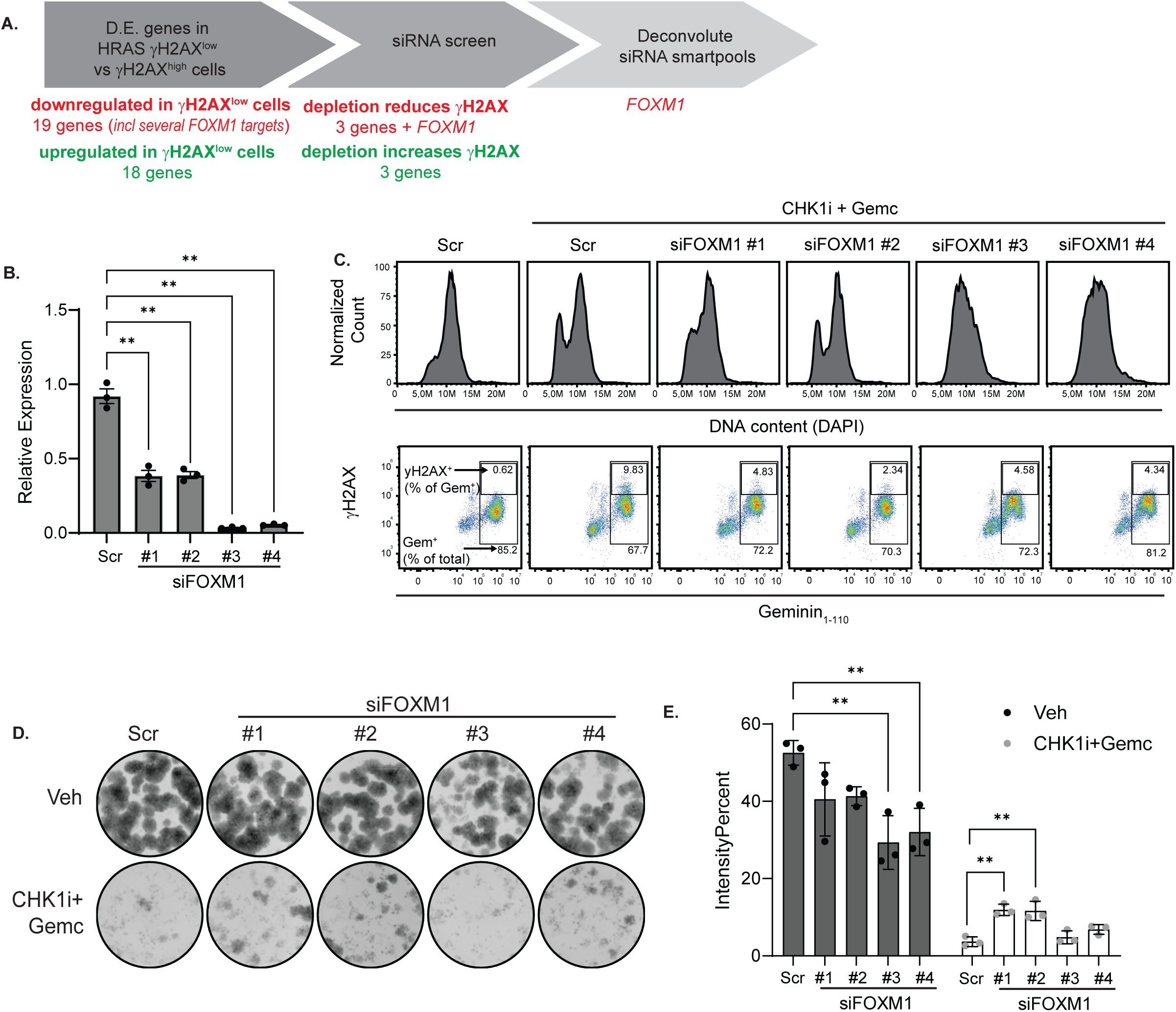
Partial knockdown of FOXM1 improves tolerance to replication stress without affecting cell proliferation. **A** Schematic representation of the experimental design to identify and validate putative RS-tolerance genes. **B** Relative *FOXM1* expression of RPE-HRAS^G12V^ following transfection with four individual siRNAs targeting *FOXM1* (1 nM each) as measured by qPCR. Gene expression was normalized to the average of two housekeeping genes (*GAPDH, 18S).* Error bars indicate mean +/-SEM. Significant differences determined by one-way ANOVA with Geisser Greenhouse correction followed by Dunnett’s multiple comparison test. **p<0.01, N = 3. Representative of 3 individual experiments. **C** Flow cytometry data of RPE-HRAS^G12V^ cells which were arrested in G1-phase after 24 hours treatment with a CDK4/6i and individual siRNAs against *FOXM1*. Subsequently, cells were released in the absence or presence of 10 nM CHK1i + 4 nM gemcitabine and harvested after 14 hours to enrich for S/G2-phase cells. DAPI staining was used to determine cell cycle progression (top row) and γH2AX staining was used to determine the degree of replication stress (bottom row). Number in bottom right corner of bottom row plots indicates the Geminin ^+^ cells as percentage of the total cells. Number in the top right corner of bottom row plots indicates γH2AX^+^ cells as a percentage of Geminin_1-110_^+^ cells. Representative of 2 individual experiments. **D** Outgrown colonies of RPE-HRAS^G12V^ cells transfected for 24 hr with siRNAs against FOXM1 followed by treatment for 48 hr with the 2 nM CHK1i and 4 nM gemcitabine. After removing drug-containing media, colonies were allowed to grow for 5 days. Error bars indicate mean +/-SEM. Representative of 2 individual experiments. **E** Quantification of colonies presented in panel B, presented as the IntensityPercent, which takes into account both the area covered by the cell growth as well as the pixel intensity of the covered area. This was quantified using the ImageJ ColonyArea plug-in. Error bars indicate mean +/-SEM. Significant differences were determined by ordinary One-way ANOVA followed by Dunnett’s multiple comparison test. *p<0.05, **p<0.01,N=3.

First, we confirmed that small interfering RNAs (siRNAs) targeting the putative RS-sensitizing mechanisms efficiently depleted their target gene (Fig. S3A). For initial siRNA knockdowns, we used siRNA Smartpools, which consist of four unique siRNAs targeting the same gene. Since RS-inducing drugs only affect replicating cells, we sought to enrich the cell population for cells in late S/G2 phase of the cell cycle at the time of analysis. To this end, we depleted the gene of interest using siRNA and arrested all cells in G1-phase using the CDK4/6 inhibitor palbociclib. Subsequently, we released the cells from palbociclib in the presence of CHK1i + gemcitabine and evaluated the level of RS 14 hours after release, when most cells were in late S/G2 phase (Fig. S3B). In addition to enriching for late S/G2 phase cells, this approach ensured that all cells start DNA replication in the presence of CHK1i + gemcitabine. To exclude bias from cells that failed to enter S-phase after palbociclib release when evaluating the level of RS, we calculated the γH2AX positive cells as percentage of the Geminin_1-110_ positive (i.e., S/G2-phase) cells.

We first evaluated if depletion of genes upregulated in γH2AX^low^ cells could resensitize cells to RS (Fig. 4A). While depletion of most putative RS-tolerance genes using Smartpool siRNAs did not affect RS, depletion of *MYL6*, *PRDX5* and *ARL4C* increased RS induced by CHK1i + gemcitabine (Fig. S3C). However, when we then used individual siRNAs to deconvolute the effects of the Smartpools, none of the three targets showed a consistent RS-sensitizing effect, suggesting off-target effects of the individual siRNAs (data not shown).

We then shifted our focus to genes that may make cells more sensitive to RS. *AMOTL2*, *CTGF* and *CYR61* were downregulated in γH2AX^low^ cells and knockdown reduced the fraction of cells with severe RS (Fig. S3C). This suggests that these genes sensitize cells to RS-inducing drugs. Similarly, knockdown of *FOXM1,* which regulates the expression of a panel of sensitizing genes (*CENPE*, *UBE2C*, *HMGB2*, *ANLN*, *MKI67*) bolstered resistance to RS induced by CHK1i + gemcitabine (Fig. SC3). However, only *FOXM1* knockdown passed the deconvolution step, i.e., consistent phenotypes with individual siRNAs.

We observed varying levels of *FOXM1* knockdown with the four siRNAs against *FOXM1* present in the siRNA Smartpool, with two siRNAs accomplishing a near-complete knockdown (<90%) and two accomplishing a modest knockdown (50-60%) of the gene (Fig. 4B). The siRNAs that produced stronger *FOXM1* knockdown also slowed cell cycle progression, as seen by a lower proportion of cells in G2 phase at 14 hours following release from G1 arrest (Fig. 4C, top row). Despite the differential degrees of *FOXM1* expression and cell cycle progression, all four knockdowns resulted in decreased levels of γH2AX relative to cells transfected with scrambled siRNA after 14 hr of CHK1i + gemcitabine treatment in synchronized RPE-HRAS^G12V^ cells (Fig. 4C, bottom row; quantified in Fig. S4).

We then performed clonogenic survival assays in cells with varying *FOXM1* gene expression levels during treatment with CHK1i + gemcitabine (Fig. 4D,E). The transient effect of siRNA knockdown allowed us to focus on the consequence of reduced FOXM1 expression at the time of treatment (mimicking stochastic variation in expression levels that may occur in cancer cells) rather than a permanent change in expression levels. We quantified colony outgrowth using the IntensityPercent value calculate by the ImageJ ColonyArea plugin, which takes into account both the area covered by cells as well as the density of the cells within a colony (Guzman et al., 2014). In untreated cells, the greater knockdown (siRNA#3 and siRNA#4) resulted in fewer colonies, which is expected since knockdown of *FOXM1* slows cell proliferation. The partial knockdown had little to no effect on colonies in untreated cells, consistent with the minimal effect on cell cycle progression. In the scrambled condition, treatment with CHK1i + gemcitabine nearly suppressed the outgrowth of colonies once the drugs were removed. However, in the partial knockdowns, several dense colonies were able to recover even following drug exposure, suggesting that the partial knockdown of FOXM1 protected cells from RS-induced cell death without limiting cell proliferation.

To summarize, these data indicate that partial FOXM1 knockdown protects against drug-induced RS while still allowing cells to proliferate. This suggests that cancer cells may use similar tuning of the FOXM1 expression program to resist the effects of RS-inducing drugs without fully halting growth.

## Discussion

In this study, we used single cell RNA sequencing to investigate transcriptional heterogeneity associated with differential responses to RS. We employed a novel technique wherein cells are reversibly fixed, allowing for cell sorting based on intracellular staining, and subsequent single cell RNA sequencing on sorted cells. By comparing cells with low levels of RS (as measured by γH2AX following treatment with RS-inducing drugs) to cells with high levels of RS, we found that a moderate reduction in gene expression of downstream targets of transcription factor *FOXM1* protects cells against RS induced by CHK1i+gemcitabine without significantly delaying cell proliferation.

Compared to popular screening approaches such as CRISPR or RNA interference libraries, our approach has caveats. First, a differentially expressed gene in RS-tolerant versus RS-sensitive cells can be either a cause or a consequence of the observed phenotype. Our observation that P53 target genes were downregulated in RS-tolerant cells illustrates this issue (Figure 3E, S2C). The far majority of our potential hits did not show consistent phenotypes when knocked down with siRNA, suggesting that they do not play a direct role in regulating RS. Second, genes may act in concert: the effect of the variation in expression of an individual gene can be minimal, while up- or down-regulation of multiple genes can have a tolerizing effect. Further studies altering the levels of multiple genes at once would be necessary to test this hypothesis. Nevertheless, our single cell transcriptomics approach has the advantage over the aforementioned screening approaches that it has the potential to detect the effects of stochastic heterogenous transcription events within the physiological range.

We observed that reducing FOXM1 expression improves cell viability (as seen by colony outgrowth) following treatment with RS-inducing drugs. This is consistent with recent studies showing that high FOXM1 activity facilitates unscheduled mitotic entry to cause RS-induced mitotic catastrophe (O’Brien et al., 2023, Gallo et al., 2022, Chung et al., 2019, Branigan et al., 2021). FOXM1 primes cells to enter mitosis by inducing the transcription of a large set of mitotic genes, including *CCNB1, PLK1* and *CDK1* itself which allows for sufficient accumulation and activation of cyclin B-CDK1 complexes to enter mitosis (Sanders et al., 2015, Sadasivam, Duan & DeCaprio, 2012). Curtailing CDK1 activation reduces sensitivity to RS response inhibitors because cells are less prone to enter mitosis prematurely with DNA damage sustained during RS. This can be reversed by inhibiting key regulators of CDK1 activity, such as WEE1, thus reactivating CDK1 (Ruiz et al., 2016).

In addition to highlighting the described role of FOXM1 in promoting mitotic catastrophe, this study also points to a RS-mediating activity of FOXM1 during S-phase. Our finding that low FOXM1 expression reduces CHK1i-mediated DNA damage is consistent with recent observations showing that FOXM1 deletion reduced replication stress and DNA damage in S-phase during CHK1i treatment (Branigan et al., 2021). Remarkably, Braningan and co-workers observed no effect of full FOXM1 deletion on cell cycle progression, whereas we observed that near-complete knockdown of FOXM1 caused a clear reduction of S phase progression and proliferation rates (Figure 4B, C; siRNAs #3 and #4). A difference could be that FOXM1-mutant cells have adapted to chronic loss of this transcription factor in the former study, where RNAi-mediated knockdown provides a more acute setting. Notwithstanding these differences, the mechanism by which high FOXM1 activity is a prerequisite to accumulate DNA damage in S-phase during CHK1 inhibition remains to be uncovered. One possible explanation could be that high FOXM1 expression triggers excessive origin firing during CHK1 inhibition. An important function of CHK1 is to mitigate DNA damage by reducing firing of late origins under conditions of replication stress (Baillie, Stirling, 2021). Cyclin A2-CDK1 complexes mediate origin firing, and *CCNA2* is a FOXM1 target (Katsuno et al., 2009), thus FOXM1-induced CCNA2 expression exacerbate the increase in late origin firing permitted by CHK1 inhibition. Consistent with this, a recent study using ATR-deficient B cells showed that RS triggered by loss of ATR could be reversed by suppressing origin firing, which was accomplished through partial inhibition of CDC7 and CDK1 activity (Menolfi et al., 2023).

The work described here describes a model in which transcriptomic variability of the transcription factor FOXM1 endows a subset of cells within a population of genetically identical cells tolerance to drug-induced replication stress. While it may be hard to therapeutically target FOXM1 to improve efficacy of intra-S-phase checkpoint inhibitors, overexpression of the FOXM1 program can potentially serve as a biomarker since amplification of the FOXM1 gene occur relatively frequently in multiple cancers (Barger et al., 2019)., although single cell analysis would need to reveal the relative heterogeneity within tumors. Furthermore, our findings support the idea that decelerated S-phase progression could counteract CHK1 inhibitors, which also suggests that pharmacologically accelerating cell cycle progression may work to sensitize cells to this class of drugs. An excellent example of this is inhibiting WEE1, the kinase responsible for preventing CDK1/2 activation or its relative PKMYT1, which inhibits CDK2. WEE1 and PKMYT1 inhibitors force cell proliferation in the presence of RS and – at least in part – overcome resistance to intra S-phase checkpoint inhibitors (Ruiz et al., 2016, Chung et al., 2019, Koh et al., 2018).

## Methods

### Key resources

Key resources are listed in Table S1.

### Cell lines

hTERT RPE-1 cell line was obtained from ATCC and cultured at 37°C, 5% CO_2_ in DMEM supplemented with 10% FBS and 1% pen/strep. Cell lines were regularly tested and confirmed mycoplasma negative.

Overexpression of HRAS^G12V^ was induced by adding 0.2 µg/mL doxycycline to the culture medium. Gemcitabine, prexasertib and palbociclib were purchased from Selleck chemicals and used at a final concentration of 4 nM, 10 nM and 1 μM respectively, unless stated otherwise.

RPE cell lines harboring the Tet Repressor, doxycycline inducible HRAS^G12V^, FUCCI4 system and fluorescent tagged truncated 53BP1 were created using the third-generation lentiviral packaging system as previously described (Segeren et al., 2022).

### DSP fixation and antibody staining of single cells

Fixation of cells was performed according to the protocol described by Gerlach *et al*. (Gerlach et al., 2019). In short, cells were collected by trypsinization, washed with PBS after which cells were fixed with 0.5mM dithiobis(succinimidyl propionate) (DSP) in Sodium Phosphate buffered Saline (pH 8.4) for 45 minutes at room temperature at a concentration of 1 million cells per 2.5mL. Next, DSP was neutralized by incubating the cells with quench buffer (100mM Tris, pH 7.5, 150mM NaCl) for 10 minutes and cell clumps were removed using a 70 μm cell strainer. Cells were incubated for 30 minutes with BP buffer (PFBB:PBS, 1:1, supplemented with 0.1% Triton X-100) to permeabilize the cells, after which samples were incubated overnight with antibodies in BP buffer. If samples were intended to use for single-cell RNA-sequencing, BP buffer was supplemented with 2 U/μl RNAsin Plus. Samples were filtered using a 40μm cell strainer and incubated with DAPI (0.2μg/mL) prior to loading of samples on the flow cytometer.

For antibody testing, samples were loaded on a CytoFLEX flow cytometer and analyzed using FlowJo v10.0 software. Index sorting of cells for single-cell RNA-sequencing was performed on a BD Influx cell sorter.

### Microscopy

For immunofluorescence staining, cells were seeded on coverslips. Prior to fixation of cells using 4% paraformaldehyde for 20 minutes, pre-extraction with 0.2% Triton X-100 for 1 minute on ice was performed. Next, cells were permeabilized using 0.1% Triton X-100 for 10 minutes, blocked with 5% goat serum and incubated with indicated antibodies after which coverslips were mounted on slides. Samples were analyzed on a Leica SP8 confocal microscope equipped with a 20x objective. Antibodies and dilutions used are listed in Table S2.

### Immunoblotting

For immunoblotting, cells were washed twice with ice-cold PBS and lysed in ice-cold RIPA-buffer (50 nM Tris-HCl pH 7.5, 1 mM EDTA, 150 mM NaCl, 0.25% deoxycholate, 1% NP-40) supplemented with NaF (1 mM), NaV_3_O_4_ (1 mM) and protease inhibitor cocktail (11873580001, Sigma Aldrich) after which samples were subjected to a standard SDS-page immunoblot. Antibodies used and dilutions are listed in Table S3.

### RNAi transfections

For siRNA experiments, cells were transfected with siRNA targeting the gene of interest or a scrambled control using Lipofectamine RNAiMAX according to manufacturers’ instructions (Life Technologies, 13778030). ON-TARGETplus SMARTpool siRNAs were purchased as custom cherry-pick libraries from Dharmacon and used at a final concentration of 10 nM, while individual siRNAs were used at a final concentration of 1 nM. Efficient knock-down of intended target was confirmed by quantitative PCR 24 hours after transfection.

### Fixation and staining to validate hits with flow cytometry

Cells were collected by trypsinization, washed with PBS, transferred to a 96 well plate and fixed using 4% PFA for 30 minutes while gently shaking. Next, cells were washed with 0.1% BSA in PBS and permeabilized using 0.1% Triton for 30 minutes. Cells were washed once more with 0.1% BSA in PBS prior to incubation with the fluorescent linked γH2AX antibody for 1 hour at room temperature. DAPI was added to the samples at a final concentration of 2.0μg/100,000 cells to stain DNA content. Samples were loaded on a CytoFLEX flow cytometer and analyzed using FlowJo v10.0 software.

### Clonogenic survival assays

Cells were seeded at low density (200 cells per 12 well plate) to assess colony formation. Following 24 hour transfection with 1 nM siRNAs targeting FOXM1, cells were treated with 2 nM prexasertib and 4 nM gemcitabine. After 48 hr exposure to the drugs, media was replaced with drug-free media and remaining cells were allowed to grow out to form colonies. The ImageJ ColonyArea plug-in was used to quantify the area of the well covered by colonies as previously described (Guzman et al., 2014).

### Quantitative PCR

RNA isolation, reverse transcription and quantitative PCR were performed according to manufacturers’ instructions using the QIAGEN RNeasy kit, Thermo Fisher cDNA synthesis kit and Bio-RAD SYBR Green Master mix, respectively. Quantification of relative gene transcript levels was performed using the ΔΔCt method for multiple-reference gene correction using GAPDH or β-Actin and RPS18 as reference genes. Primers used in this manuscript are listed in Table S3.

### Single-cell RNA-sequencing

For single-cell RNA-sequencing single cells were collected in 384-well plates containing barcoded CEL-seq2 primers and nucleotides using index sorting and stored at −80°C until further processing. De-crosslinking of the DSP fixed cells was performed by addition of 0.1M DTT to the reverse transcription mix (10 mM DTT final concentration), which is part of the regular reverse transcription mix for unfixed cells. SORT-seq sequencing and read alignment were performed by Single Cell Discoveries (Utrecht, the Netherlands) using their pipeline based on CEL-Seq2 (Muraro et al., 2016, Hashimshony et al., 2016). Briefly, samples were barcoded with CEL-seq2 barcodes and UMI during reverse transcription and pooled after second strand synthesis. The resulting cDNA was amplified with an overnight in vitro transcription reaction. From this amplified RNA, sequencing libraries were prepared with Illumina TruSeq small RNA primers, which were paired-end sequencing on the Illumina NextSeq500 platform. Read 1 was used to identify the Illumina library index and CEL-seq2 sample barcode. Read 2 was aligned to the human genome (hg38) transcriptome using the Burrows–Wheeler Aligner v0.7.17. Reads that mapped equally well to multiple locations were discarded. Mapping and generation of count tables were done using the MapAndGo2 script. Downstream processing and analysis were performed in Rstudio (Version 1.4.1106) and R (Version 4.0.5) using the Seurat package (Version 3.2.3) (Stuart et al., 2019). Cells were filtered and selected for downstream analysis when the following parameters were met: number of detected genes > 1000 and < 10,000, Unique Molecular Identifier (UMI) counts > 3,000 and < 75,000, and the percentage of mitochondrial counts and ERCC RNA spike-ins below 25 and 5 respectively. Next, raw counts were normalized, and variance stabilized using the SCTransform method (Hafemeister, Satija, 2019). Subsequently, dimension reduction was performed by principal component analysis. Identified clusters were visualized with t-Distributed Stochastic Neighbor Embedding (t-SNE). Differentially expressed genes were identified using the Seurat FindAllMarkers() function with one non default argument, min.pct = 0.25 requiring a greater fraction of cells within a cluster to have expression. After this the results were filtered at a Bonferroni adjusted significance level of p < 0.05. Expression correlation between the differentially expressed genes was determined using Pearson correlation. All sequencing data generated in this study are available on the Gene Expression Omnibus under accession numbers GSE256134 and GSE250285.

### Quantification and statistical analysis

Flow cytometry, immunoblot and quantitative PCR experiments were performed three times unless indicated otherwise. Details on sample size and statistical methods employed are described in the figure legends. * p < 0.05, ** p < 0.01, *** p < 0.001 unless indicated otherwise.

## Acknowledgements

We thank Reinier van der Linden and Stefan van der Elst (Hubrecht Institute-KNAW, NL) for assistance with FACS sorting experiments. We thank Klaas Mulder and Jan Gerlach for inspiring us to establish DSP fixation for single cell RNA-sequencing, and for helpful practical suggestions. We thank Utrecht Sequencing Facility for providing sequencing service and data. This work is financially supported by the KWF Kankerbestrijding (Dutch Cancer Society, project grant 11941-2018-II) and ZonMW (grant 91116011). Further financial support was provided by research infrastructure grants from Utrecht Life Sciences to the Single Cell Analysis Center and the Center for Cell Imaging. Utrecht Sequencing Facility is subsidized by the University Medical Center Utrecht, Hubrecht Institute, Utrecht University and The Netherlands X-omics Initiative (NWO project 184.034.019). We thank Alain de Bruin and the other members of the Cancer Group for constructive feedback and suggestions.

## Author Contributions

H.A.S. conceived and performed experiments, analyzed data, and wrote the manuscript. K.A.W. conceived and performed experiments, analyzed data, and wrote the manuscript. E.A.v.L performed experiments and analyzed data. F.M.R. analyzed single cell RNA-sequencing data. B.W. conceived and oversaw the study and wrote the manuscript.

## Competing Interests

The authors declare no competing interests.

## Data Availability

All sequencing data is available at GEO under accession number GSE256134 for fresh and RAID-fixed cells (Figure 2) and accession number GSE250285 for RAID-fixed cells sorted by yH2AX level (Figure 3).

## Notes

https://www.ncbi.nlm.nih.gov/geo/query/acc.cgi?acc=GSE256134

https://www.ncbi.nlm.nih.gov/geo/query/acc.cgi?acc=GSE250285

### Competing Interest Statement

The authors have declared no competing interest.

