## Supplementary figures and images for "Stochastic variation in the FOXM1 transcription program mediates replication stress tolerance"

### Supplemental Figure 1

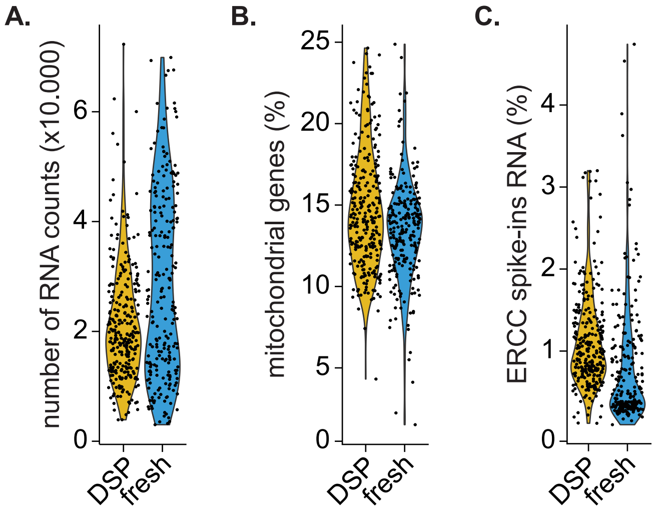


**Supplemental Figure 1, related to figure 2**

### Supplemental Figure 2

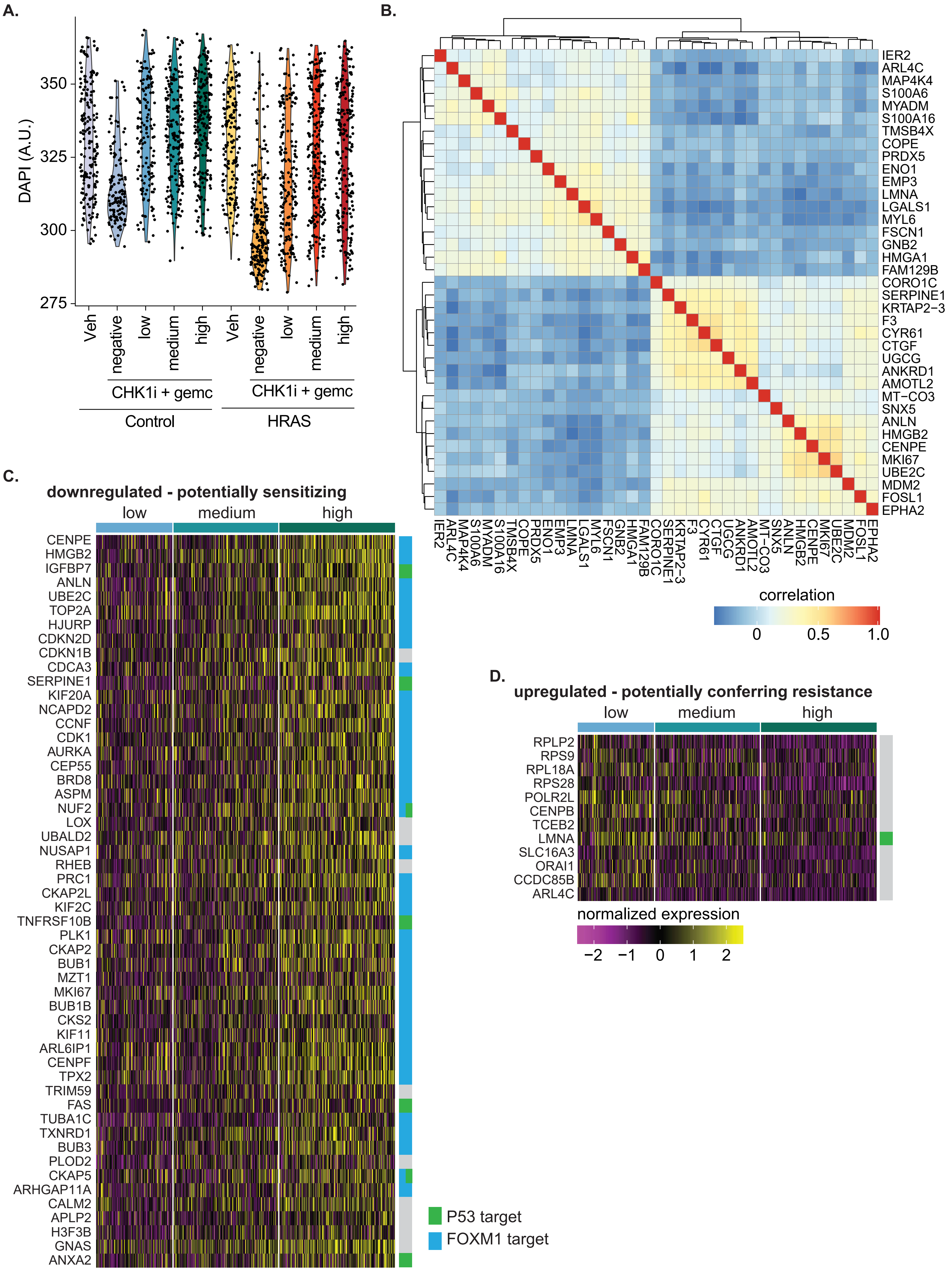


**Supplemental Figure 2, related to figure 3**

### Supplemental Figure 3

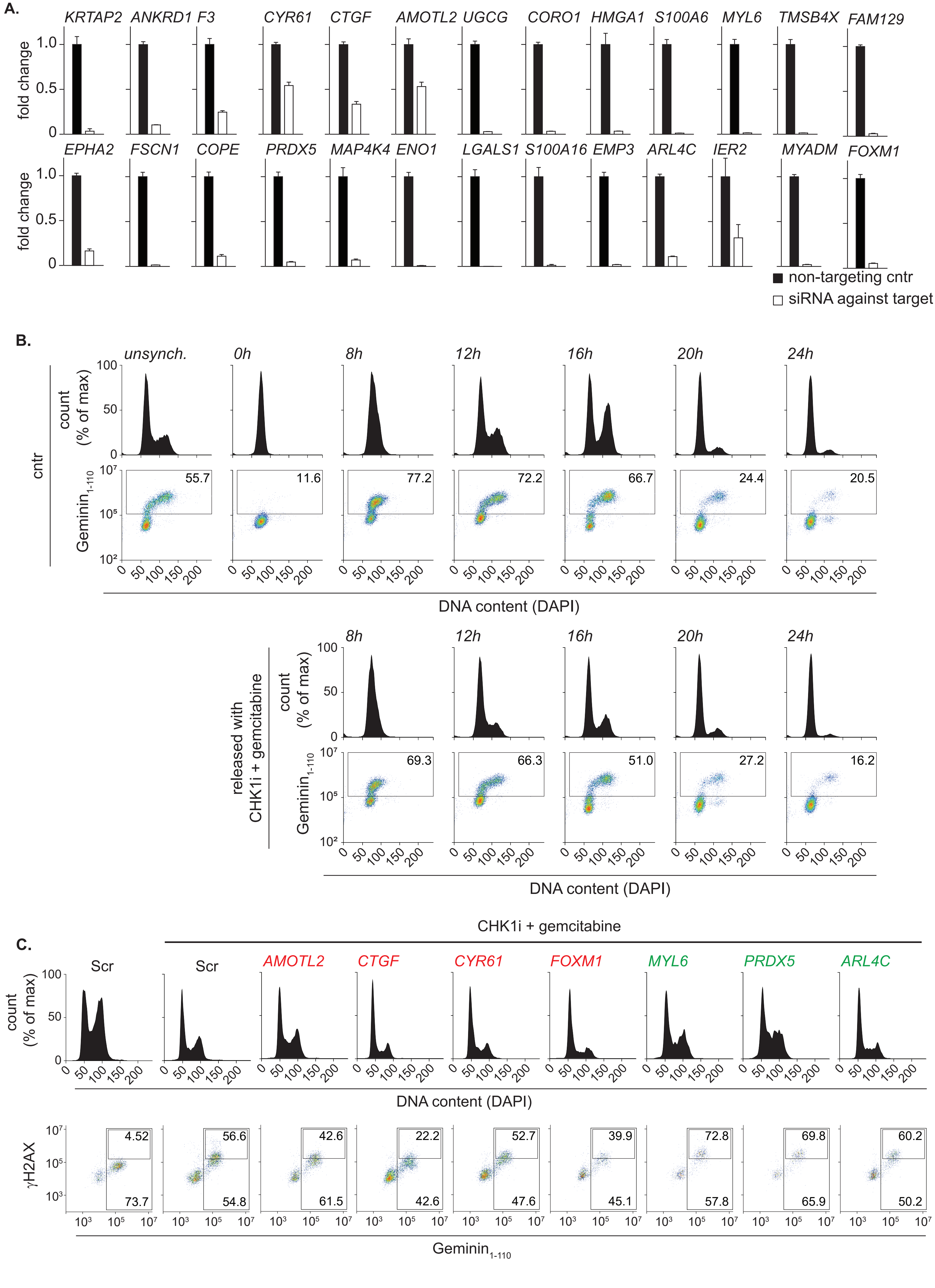


**Supplemental Figure 3, related to figure 4A**

### Supplemental Figure 4

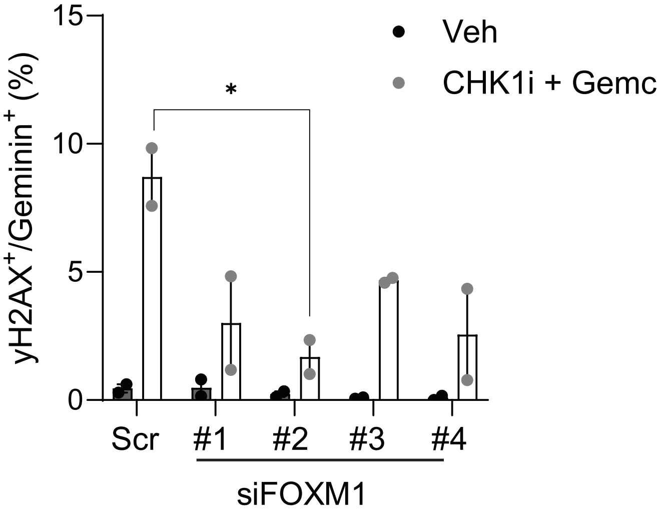


**Supplemental Figure S4, related to figure 4C**
