## Supplemental Information for "Stochastic variation in the FOXM1 transcription program mediates replication stress tolerance"

### 1. Supplementary Tables

**Table S1. Key resources**

| **Reagent or resource** | **Source** | **Identifier** |
|---|---|---|
| **Antibodies** | | |
| CHK1 phospho S296 | Cell Signaling Technology | 2349; RRID:AB_2080323 |
| CHK1 phospho S345 | Cell Signaling Technology | 2348; RRID:AB_331212 |
| CHK1 | Cell Signaling Technology | 2360; RRID:AB_2080320 |
| γ-H2AX (S139) | Cell Signaling Technology | 2577; RRID:AB_2118010 |
| γ-TUBULIN | Sigma-Aldrich | T6557; RRID:AB_477584 |
| CHK1 phospho S345 - PE conjugated | Abcam | Ab278744; RRID:AB_2745001 |
| KAP1 phospho S824 | Bethyl | A300-767A; RRID: AB_669740 |
| RPA2 phospho S8 | Cell Signaling Technology | 54762S; RRID: AB_2799471 |
| γH2AX – Alexa647 conjugated | BioLegend | 613407; RRID: AB_2114994 |
| **Chemicals** | | |
| Palbociclib | Selleck chemicals | S1116 |
| Gemcitabine | Selleck chemicals | S1714 |
| Prexasertib | Selleck chemicals | S7178 |
| DAPI | Sigma-Aldrich | D9542 |
| Protease inhibitor Cocktail | Sigma-Aldrich | 11873580001 |
| Fetal bovine serum | Thermo Fisher | 10500064 |
| DMEM | Thermo Fisher | 41966052 |
| **Critical commercial assays** | | |
| Rneasy Micro Kit for RNA extraction | Qiagen | 74004 |
| **Deposited data** | | |
| Single-cell RNA sequencing RPE-FUCCI4 HRAS$^{G12V}$ cells | This paper | |
| **Experimental Models** | | |
| hTERT RPE-1 cells | ATCC | CRL-4000; RRID:CVCL_4388 |
| **Oligonucleotides** | | |

| | | |
|---|---|---|
| Primers used for qPCR, see Table S3 | This paper | Biolegio |
| Scrambled siRNA | Dharmacon | D-001210-02-05 |
| **Software and Algorithms** | | |
| FIJI (ImageJ) | https://fiji.sc | RRID:SCR_002285 |
| FlowJo | BD | RRID:SCR_008520 |
| R | https://www.R-project.org/ | N/A |
| Rstudio | https://www.rstudio.com/ | RRID:SCR_000432 |
| USEQ RNA-seq pipeline | https://github.com/UMCUGenetics/RNASeq | N/A |

**Table S2. Antibodies for immunoblots and immunofluorescence staining**

| *Application* | *Name* | *Company* | *Catalogue number* | *Dilution* |
|---|---|---|---|---|
| *Immunoblots* | CHK1 phospho S296 | Cell Signaling | 2349 | 1:1000 |
| | CHK1 phospho S345 | Cell Signaling | 2348 | 1:1000 |
| | CHK1 | Cell Signaling | 2360 | 1:1000 |
| | KAP1 phospho S824 | Bethyl | A300-767A | 1:500 |
| | γ-H2AX (S139) | Cell Signaling | 2577 | 1:1000 |
| | γ-TUBULIN | Sigma-Aldrich | T6557 (GTU-88) | 1:1000 |
| *Immunofluorescence* | γ-H2AX (S139) | Cell Signaling | 2577 | 1:200 |
| *Flow cytometry* | CHK1 phospho S345 – PE conjugated | Abcam | Ab278744 | 1:5000 |
| | KAP1 phospho S824 | Bethyl | A300-767A | 1:500 |
| | RPA2 phospho S8 | Cell Signaling | 54762S | 1:500 |
| | γH2AX – Alexa647 conjugated | BioLegend | 613407 | 1:200 |

**Table S3. qPCR primers**

| **Gene** | **Forward primer (5’-3’)** | **Reverse primer (3’-5’)** |
|---|---|---|
| *AMOTL2* | CTACAGCAGACAGAGCACCC | ATCTCTGCTCCCGTGTTTGG |
| *ANKRD1* | CGGAGCATCTTATCGCCTGT | TTCTGCCAGTGTAGCACCAG |
| *ARL4C* | TTGGTTCGCTCTTTGTTCGC | GAAACGCAGGAAGTCCCTCA |
| *β-ACTIN* | GATCGGCGGCTCCATCCTG | GACTCGTCATACTCCTGCTTGC |
| *COPE* | GCCATGACAGTGCAGATCCT | GCACTTGTCAGCCATCTCCT |
| *CORO1* | TTTGCAGCGGGGACTTCG | TTGCTCTGTCGTACCACTCG |
| *CTGF* | ATGGTGCTCCCTGCATCTTC | CTGGTACTTGCAGCTGCTCT |
| *CYR61* | CCAGTGTACAGCAGCCTGAA | CGCATCTTCACAGTCCTGGT |
| *ENO1* | GGCTGTTGAGCACATCAATAAAAC | GCACCAGCTTTGCAGACG |
| *EMP3* | CGAGGGACAAGACTCCGAC | TTGTCCAAAGTGGCCACGAA |
| *EPHA2* | TCACACACCCGTATGGCAAA | ACGTTGCACACGGAGTACAT |
| *F3* | CAGCCCGGTAGAGTGTATGG | AGCTCCAACAGTGCTTCCTT |
| *FAM129B* | CCCTTCCTTTTGGGGCTCTC | AAGAGAGCCACGCCATACTG |

| | | |
|---|---|---|
| *FOXM1* | AGACCTGTGCAGATGGTGAG | CTGATGGTCTCGAAGGCTCC |
| *FSCN1* | AAGGACGAGCTCTTTGCTCT | CGGTCTCCTCGTCCTGATTG |
| *GAPDH* | CTCTGCTCCTCCTGTTCG | GCCCAATACGACCAAATCC |
| *GNB2* | ACAGTGGGTTTTGCTGGACA | CGTAGCCGTTGGGGAAGAAA |
| *IER2* | GTGTCGGAGTTCTGTCTGGG | CCAAACACTCATTGCCCGTG |
| *LGALS1* | TCGGGTGGAGTCTTCTGACA | CAGGTTCAGCACGAAGCTCT |
| *LMNA* | GCGTACGGCTCTCATCAACT | CGAGCGCAGGTTGTACTCA |
| *HMGA1* | GCTCCTCTAATTGGGACTCCG | TCCTTTTCCTGCTTGGAGGC |
| *MAP4K4* | TTTTCAGACCCCTCAAGCCT | TTGTGAGGTGGCCGTACATC |
| *MYADM* | ACAGCCTGTTCCAAGTGTGG | AAGATGACGTCGTGGTGGTT |
| *MYL6* | GTCGAAGGACTTCGGGTGTT | AAGGTCCTCAGCCATTCAGC |
| *KRTAP2-3* | AGCTGATCCTCAAGCACGAA | GGGTGATGAGTCAGTGGGAC |
| *PRDX5* | ACGCTCAGCGGGCTATATACT | TCAAACACCTCCACTGCTGG |
| *RSP18* | AGTTCCAGCATATTTTGCGAG | CTCTTGGTGAGGTCAATGTC |
| *S100A16* | CCAAATCTGACTGTGGCTTGC | CTGACATCTCCCTGCTTCGC |
| *S100A6* | ATTTGGCCGCCTCCCTACC | TTCTTGCTCAGGGTGTGCTT |
| *TMSB4X* | CCAGACTTCGCTCGTACTCG | CGCACGCCTCATTACGATTC |
| *UGCG* | TTGCTGCCACCTTAGAGCAG | TCGGTCAGCTATCGCTTTGG |

### 2. Supplementary Figure Legends

**Figure S1. Related to Figure 2**

**A** Violin plot showing the number of unique RNA (UMI) counts per cell in DSP-fixed and fresh cells RPE-$HRAS^{G12V}$ cells.

**B** Violin plot showing the percentage of RNA counts mapping to mitochondrial genes as percentage of total counts detected in DSP-fixed and fresh cells RPE-$HRAS^{G12V}$ cells.

**C** Violin plot showing ERCC spike-in RNA counts as percentage of total counts detected in DSP-fixed and fresh RPE-$HRAS^{G12V}$ cells.

**Figure S2. Related to Figure 3**

**A** The DAPI fluorescence intensity in individual FACS-sorted cells shown in a violin plot. Note that only cells negative for γH2AX display substantial lower DAPI levels. The lower DAPI signal in RPE-$HRAS^{G12V}$ cells can be attributed to the higher proliferation rate, and subsequent lower DAPI to cell ratio.

**B** Correlation matrix displaying the correlation between all the differentially expressed genes in RPE $HRAS^{G12V}$ $\gamma H2AX^{high}$ versus $\gamma H2AX^{low}$ cells.

**C** Heatmap of genes differentially expressed and downregulated in $\gamma H2AX^{low}$ versus $\gamma H2AX^{high}$ control RPE cells after treatment with CHK1i + gemcitabine.

**D** Heatmap of genes differentially expressed and upregulated in $\gamma H2AX^{low}$ versus $\gamma H2AX^{high}$ control RPE cells after treatment with CHK1i + gemcitabine.

**Figure S3. Related to Figure 4A**

**A** Quantitative PCR of the expression of potential RS-tolerance conferring genes in RPE-$HRAS^{G12V}$ cells treated with scrambled siRNA or siRNA targeting the gene of interest. Gene expression was normalized to the average of two housekeeping genes (*GAPDH, 18S).* Bar represents mean ± s.e.m.

**B** Flow cytometry data of RPE-$HRAS^{G12V}$ cells unsynchronized, arrested in G1-phase after 24 hours treatment with a CDK4/6i and at indicated hours after release in the presence and absence of CHK1i + gemcitabine to enrich for S/G2-phase cells. Representative of 2 independent experiments.

**C** Flow cytometry data of RPE-$HRAS^{G12V}$ cells with the indicated genes depleted by Smartpools of four individual siRNAs. Representative of 2 independent experiments.

**Figure S4. Related to Figure 4C**

Quantification of percent of γH2AX-positive cells in geminin-positive cells (representative of cells in S/G2 phase) from two individual experiments. Error bars indicate mean +/- SEM. *$p<0.01$.
